# Subjective assessment and taste strips testing of gustatory function, at home, and in the lab

**DOI:** 10.1101/2022.09.11.507407

**Authors:** Tomer Green, Anne Wolf, Anna Oleszkiewicz, Anna Aronis, Thomas Hummel, Marta Y Pepino, Masha Y Niv

**Author notes:** **Correspondence to**: Prof. Masha Niv, Institute of Food Science, Biochemistry and Nutrition, The Hebrew University of Jerusalem, Herzl Street 229, Rehovot, Israel, postal code: 7610001.

## Abstract

Gustatory ability is an important marker of health status, including COVID-19 disease. We compare self-reporting with home and lab psychophysical “taste strips” tests in healthy subjects. The taste test consisted of paper strips impregnated with sweet, bitter, salty, or sour tastants, and with the trigeminal stimulus capsaicin, each in high and in low concentration. The test was carried out either in a controlled lab environment (74 participants, 47 women) with the strips being administered by the experimenter or self-administered by the participants at home (77 participants, 59 women). After self-reporting their subjective assessment of chemosensory ability, the participant identified the taste of each strip and rated intensity and pleasantness.

Identification score, intensity, and pleasantness averaged over the 8 taste strips were similar between the lab and the home-administered tests. Self-rated taste ability did not correlate with any of these scores, but strongly correlated with self-rated smell ability in the lab group (r=0.73), and moderately correlated in the home group (r=0.51). Taste identification correlated with intensity ratings (r=0.63 lab, r=0.36 home) but not with the pleasantness ratings (r=-0.14 lab, r=0.1 home).

The results of the taste strips test were similar in the lab and at home for healthy young participants and provide a baseline against which taste tests can be compared in future applications.

## Introduction

Chemical senses are important markers of health [1–3]. Decline or loss of smell and taste commonly occurs due to aging [4], chemo and radiotherapy treatments [5, 6], end-stage kidney disease [7], as side-effects of drugs [8, 9], or after viral infections of the upper respiratory [10] including infections by SARS-CoV-2 virus which cause coronavirus disease (COVID-19) [11]. Recently it has been shown that smell and taste functions are useful for monitoring COVID-19 status and recovery [12–14]. While taste assessment studies are less common than those on smell, it has been well established that taste loss is a symptom of COVID-19 [15–17], although mechanisms of taste impairment in COVID-19 are not well understood [16]. Self-reported taste changes were shown to be successful predictors of COVID-19 status for pre-omicron variants [15, 18]. In terms of psychophysical tests, screening with taste solutions and “taste strips” (taste-impregnated filter paper) were suggested to be useful methods to identify COVID-19-related taste loss [7, 19].

There are several taste tests in the field that are used to assessed taste function. Among them are evaluations of basic taste solutions using a whole mouth procedures, such as that described in the NIH toolbox in which different basic tastant liquid samples are used to assess taste intensity perception [20–22]. Detection threshold test, in which liquid solutions or filter paper disk soaked with different basic taste solutions are used to assess the smallest concentration that can be detected [23]. Other types of detection thresholds use a current directly applied to the tongue that elicits a metallic or sour sensation to assess an electric gustatory threshold (i.e., electrogustometry) [24, 25],; and “taste strips” – filter paper strips impregnated with different concentrations of tastants that are commonly used for taste identification tests [26]. Some of these methods are relatively time-consuming, ranging from 15 to 40 minutes per session, while others, such as liquid whole mouth test of quinine and of salt, require preparation time and are not easily transported [27].

Different taste assessment methods are weakly correlated with each other, except for a high correlation between filter paper disks and electrogustometry scores for the anterior two-thirds of the tongue [27].

The present study aimed to examine an easy and quick version of a taste strip test, both at home and in laboratory settings, and its correlation with the even simpler subjective questionnaires.

## Materials and Methods

### Participants

One hundred twenty-eight healthy Israelis (aged 20 to 46 years, details in Table 1) were recruited to perform the test either in an assisted setting in the lab or in an unassisted setting at home.

**Table 1.** Subjects participating in the study

| Cohort | Number of participants | Mean age (±SD) in years | Women | Men |
| --- | --- | --- | --- | --- |
| Lab-tested | 74 (49 from group 1, 25 from group 3) | 27.4 (±5.45) | 47 (63.52%) | 27 (36.48%) |
| Home tested | 77 (54 from group 2, 23 from group 3) | 25.14 (±3.42) | 59 (76.62%) | 18 (23.37%) |

Three groups of participants were tested: Group 1 - lab condition (n=49) tested between June and July 2021.

Group 2 were tested at home (n=54) between October and December 2021, with a subset (n=36) of participants which completed the test twice with interval of 1 to 45 days between tests. Group 3 participants completed both the test in the lab (n=25) and at home on the following day (n=23, two participants had technical difficulties completing the test at home), during July 2022.

Exclusion criteria consisted of pregnancy, allergy to any food or drug, hypo/hyperglycemia, kidney failure, and heart or blood pressure problems.

### Questionnaire

Before completing the sensory test, all participants responded to a questionnaire [28]. The questionnaire inquired for general information, medical history (ex. Previous surgeries, allergy, etc.), as well as self-rated ability to taste and smell using a Likert-type scale ranging from 1 – no ability to 7 - excellent ability. Twenty-four of the 128 participants completed both the lab and the home test and were also asked to rate their ability to sense specific taste modalities on a 1-7 scale.

### Taste tests

Next, the participants performed a taste strips identification test including four tastants (bitter, sweet, salty, or sour) and a trigeminally active compound (capsaicin) at high and at low concentrations, and rated intensity and pleasantness for each taste strip from 1 to 5 (Table 3).

The taste strips were impregnated with the concentrations listed in Table 2. The lab group also tasted astringent, which was later excluded due to its brown color which could bias the participants’ perception. Strips were prepared by the Hummel lab in Technische Universität Dresden.

**Table 2.** Taste strips impregnated with compounds and concentrations

| Strip impregnation |  | Gustatory |  |  |  | Trigeminal |  |
| --- | --- | --- | --- | --- | --- | --- | --- |
|  |  | Sweet | Bitter | Sour | Salty | Spicy | Astringent |
| Compound |  | Sucrose (g/ml) | Quinine hydrochloride (g/ml) | Citric acid (g/ml) | Sodium chloride (g/ml) | Capsaicin (g/ml) | Tannin (g/ml) |
| Concentration | Low | 0.05 | 0.0004 | 0.05 | 0.016 | $247 \times 10^{-5}$ | 0.1 |
| | High | 0.4 | 0.006 | 0.3 | 0.25 | $247 \times 10^{-4}$ | 0.2 |

While completing the taste test, lab group participants had their eyes closed while the experimenter placed the strip at the center of the anterior two-thirds of the tongue, as it is a highly innervated, easily accessible, and frequently affected by gustatory impairments [26]. Subjects were asked to keep the strip in their mouth for several seconds, then they were allowed to freely move their tongue and answered whether they sensed a taste. If so, subjects were asked to identify the taste, and rate its intensity and pleasantness according to the scales shown in Table 3. The tested strips were visually identical and presented in the same order and numbered with random 3-digit codes. The order was sour, sweet, bitter, and then salty, starting with low concentrations of the gustatory impregnated strips, then their higher concentrations. Trigeminal stimuli were presented last to avoid potential impairment in participant’s ability to taste the other strips as they have a persistent sensation.

**Table 3:** Intensity and pleasantness scales

| Score | Rate the **intensity** of taste/sensation: | Rate the **pleasantness** of taste/sensation: |
| --- | --- | --- |
| 5 | Very intense | Very pleasant |
| 4 | Intense | Pleasant |
| 3 | Medium | Neutral |
| 2 | Low | Not pleasant |
| 1 | Very low | Very unpleasant |
| 0 | No intensity | No score |

The home group self-administered the strips, presented in the same order

For each correctly identified basic taste strip (sweet, bitter, sour, and salty, high and low concentration each) participant received one point, resulting in a maximum score of 8 points for the basic tastes. Capsaicin score (0, 1, or 2 points) was calculated separately.

Both average intensity and average pleasantness were calculated for the four basic taste modalities (i.e., excluding capsaicin). For average intensity, if the participant has not identified the strip, 0 intensity was used as intensity score. For averaged pleasantness, the score was calculated only for strips for which a participant sensed a taste. For example, if the participant has felt any taste for 3 strips, then the score will be the sum of the pleasantness of these strips, divided by three.

### Data collection

Data were collected via “Compusense Cloud” (Compusense Inc., Guelph, Ontario, Canada), the lab group answered directly onto a desktop computer in a minimal distraction designated room. The home participants received a personal link to the questionnaire and could perform the test at the time and location of their choice. Questions and instructions appeared on the screen in Hebrew for both groups. For the lab group, the experimenter described the instructions before completing the test. While extracting the data for analysis, the identifying username was stripped to provide anonymity to the participant. Timestamps of each of the participant’s responses were recorded for each question to analyze the time of completion of the sensory section.

### Statistical analysis

Data were analyzed using python, version 3.6.9 in Google Colab Environment. For statistical tests, the SciPy package was used with version 1.8.0. Values are presented as the mean ± standard deviation (±SD), and continuous data were tested using unpaired two-tailed student’s t-test between the two cohorts and paired two-tailed student’s t-test between the two repetitions of group 2.

Pearson’s r was used to calculate correlations, while comparing between correlations Fisher r-to-z transformation depicting the z and two-tailed p-value was used. The threshold for statistical significance was set to p <0.05. Bonferroni correction was applied when comparing taste identification for each taste strip, 8 taste strips were compared hence corrected p-value was set to 0.00625 due to Bonferroni correction.

### Ethical approval

The study was approved by the Ethical Committee for the Use of Human Subjects in Research of the Hebrew University of Jerusalem (Ethical committee file number: 5.21). All participants were instructed on the experimental procedures and signed the informed consent form upon their recruitment. The lab group completed the test between June 2021 and August 2022, whereas the home group performed the test between October 2021 and August 2022.

## Results

Taste identification (ID), pleasantness, and intensity scores were compared between the lab and home groups. Identification score distributions between the two cohorts were similar. The mean taste ID score was 5.42 [±1.78] for the home group and 5.7 [±1.21] for the lab group, p=0.26. Intensity and pleasantness distributions were also similar for home and lab tests (Figure 1) with an average intensity of 2.8 [±0.62] for the lab group and 2. 82 [±0.7] for the home group, p=0.86; and an average pleasantness of 2.87 [±0.43] for lab group and 2.84 [±0.46] for the home group, p=0.7.

**Fig 1.**
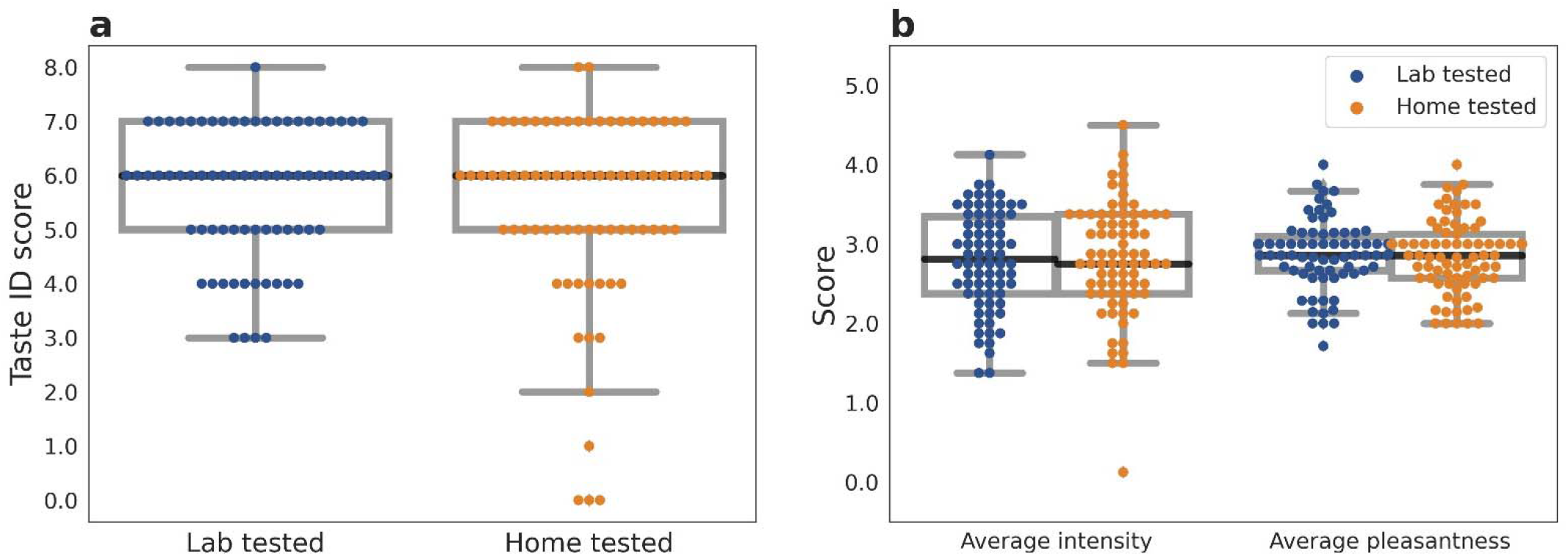
Taste identification and characteristics among the lab and home group participants. Panel A: Taste identification score distribution. Panel B: Average intensity and pleasantness ratings. Paired box plot displays the distributions of taste identification score (Taste ID), average intensity, and pleasantness scores over the 8 strips. Comparing lab and home groups colored in blue and orange respectively, the black line indicates the median, while the gray box indicates the quartiles of the dataset.

Next, we compared the correct identification of each taste at both concentrations between the two cohorts. For the low concentrations, a significant difference between the home and the lab test was demonstrated only for the spicy “low” strips, where the lab group identified the strips more accurately (Fig 2). The remaining strips at low concentration were similarly identified by both cohorts.

**Fig. 2.**
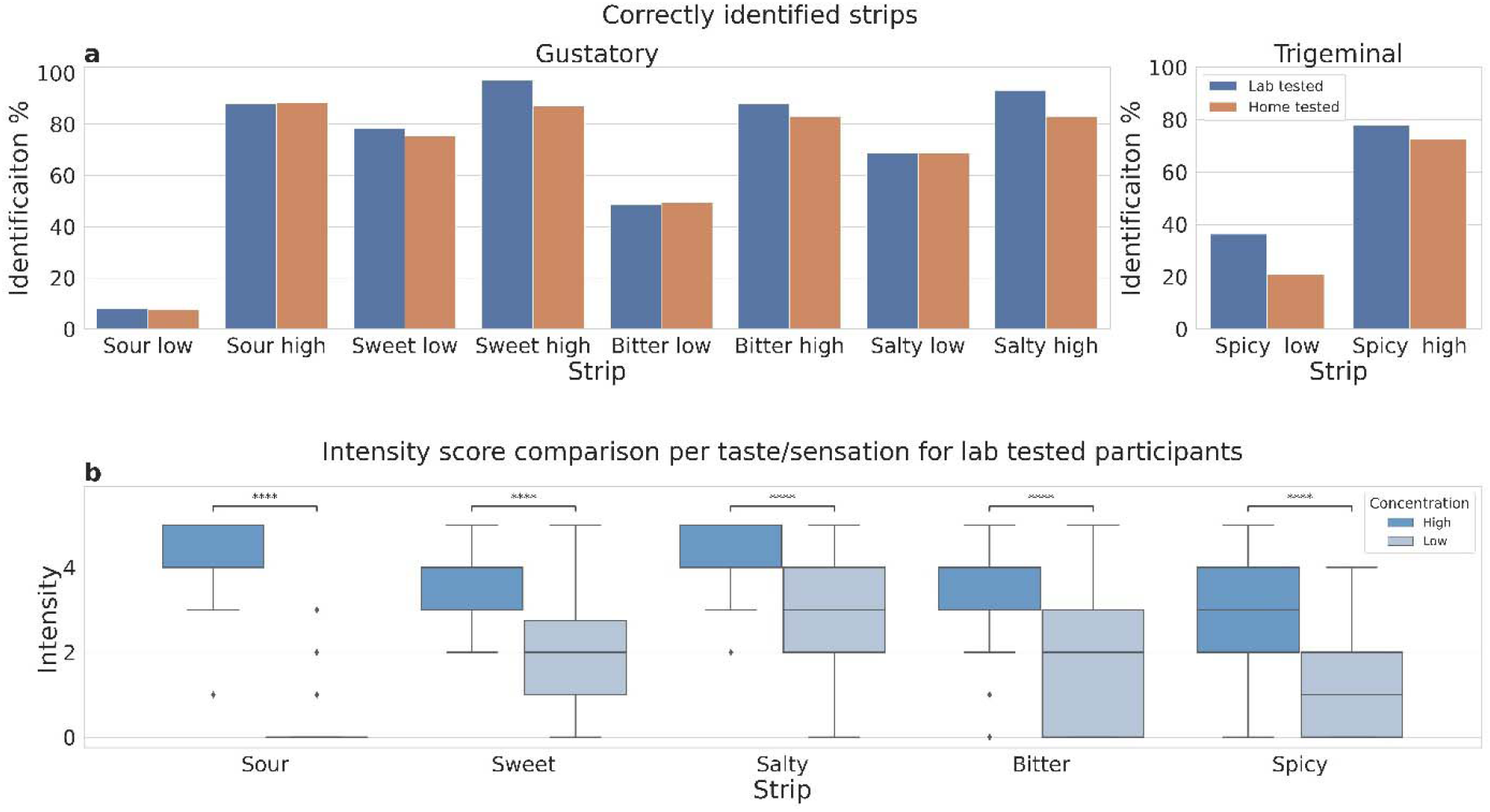
Panel A. Percent of correctly identified strips, blue for lab and orange for home testing. Panel B Intensity of each strip comparing high against low concentrations for participants tested in the lab. Independent t-test significant difference indicated by ****: p-value < 1×10^−4^.

The low concentration sour taste strip identification was the lowest in both groups, (8.1% for lab and 7.8% for home group) (Figure 2 Panel A). The sweet strip was the best identifiable strip among the low concentration strips (78.4% for the lab and 75.3% for home groups).

For the high concentration taste strips, high identification levels were observed among the lab group participants (Fig 2 A). The home cohort also identified the high concentrations well, with the exception of capsaicin, which demonstrated a small difference from lab-tested cohort (77.9% for lab and 72.7% for home setting, p=0.021).

As expected, the intensity scores were significantly lower for the low concentration strips (shown in Figure 2B for the lab group). The low sour was rarely identified (and in these cases the intensity was set to zero).

Importantly, the time did not significantly differ between the home and the lab test, as seen in Figure 3. Home tested average time to complete the sensory taste test was 8 minutes and 40 seconds [±3:28], while the time required for the lab test was slightly but not significantly longer at 8 minutes and 45 seconds [±2:05] (p=0.87).

**Fig. 3.**
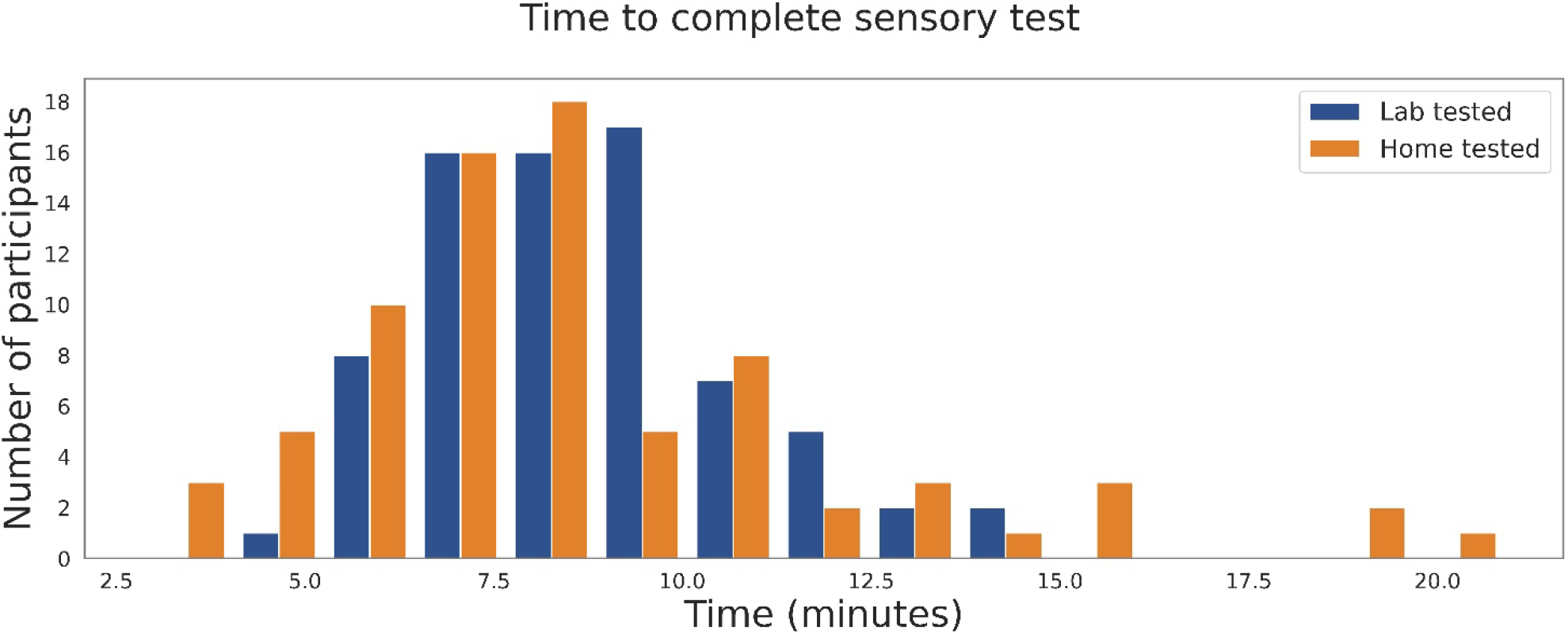
Distribution of time to complete the sensory test, lab test in blue, home test colored in orange.

Test-retest consistency of the home group (in group 2) score was assessed. The mean of the first scores deducted by the repetition scores was −0.29 (± 2.3), where almost all participants fell within one standard deviation interval as shown in Supplementary Fig 1.

Correlation analyses of self-reported parameters and objective measurements were performed, demonstrating that self-reported taste ability was strongly correlated with self-reported smell ability for the lab group (Figure 4), and somewhat correlated for the home group (Suppl. Figure 3) (r=0.73 and r=0.51, respectively, significant difference between the correlations (z = 2.2, p=0.014)). In contrast, subjective taste ability had weak or negligible correlations with taste ID, intensity ratings, or pleasantness ratings, for both the lab and the home group. Taste identification score correlated with average intensity in lab (r=0.63) but less so at home (r=0.28). It also correlated more with average pleasantness in the lab compared to the home group (r=0.48 and r=0.16, respectively).

**Fig. 4.**
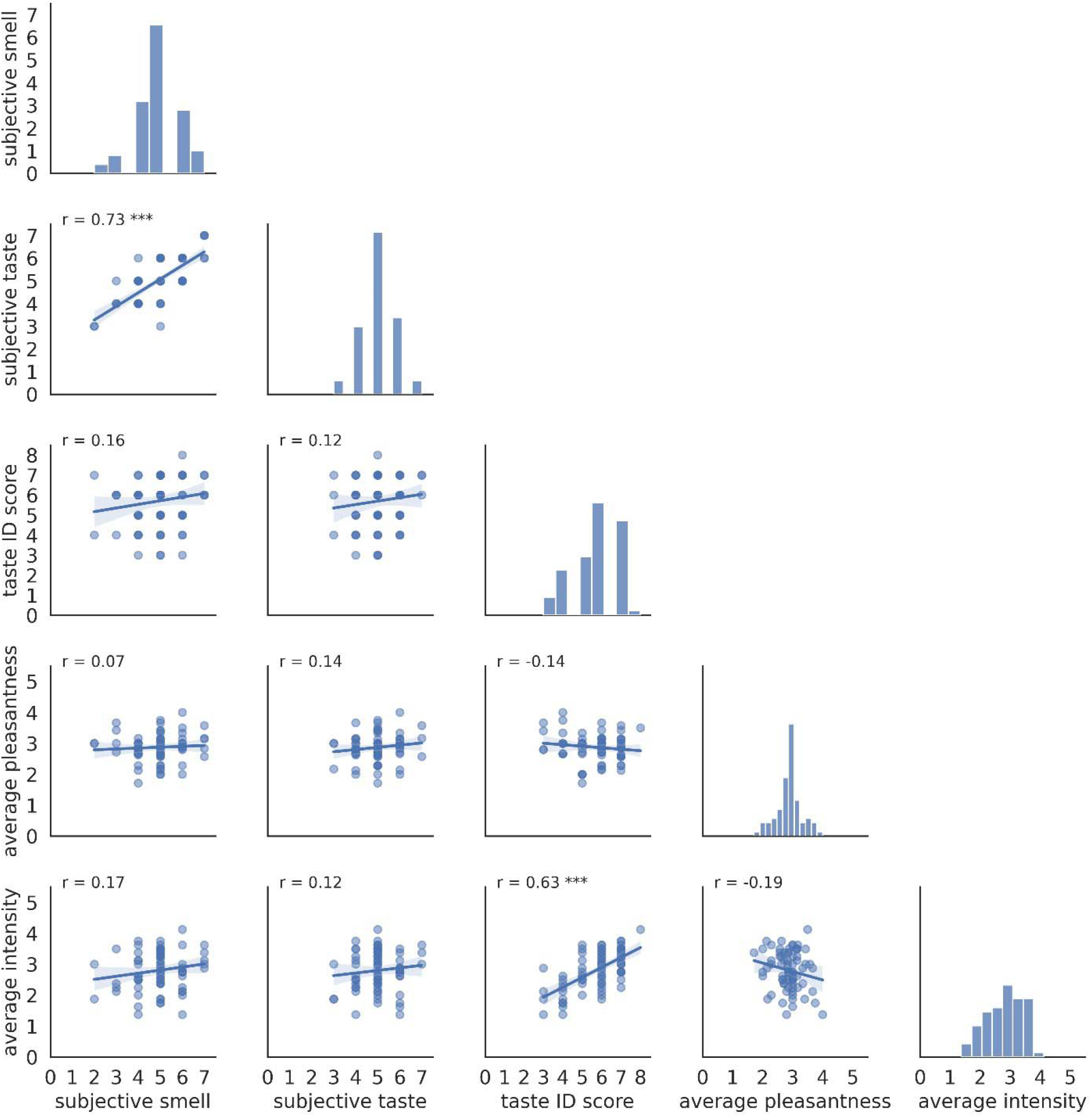
Correlations between the self-rated abilities to smell and taste (subjective smell and taste), taste identification score (taste ID score), and averaged across the four basic taste modalities regarding intensity and pleasantness ratings individually (average pleasantness and intensity), represented by scatter plots including Pearson’s R correlation coefficient at the top left of each panel, and regression line. Histograms of the diagonal panels of the figure present the frequency of scores, with the x-axis indicating the score and the y indicating the count of participants. Shown for lab group.

Participants in group 3 were asked to rank subjective ability for individual taste modalities. We found only weak correlations between subjective ability to taste the taste modality and the actual identification and intensity scores for that modality. The highest correlation was found for the subjective ability to sense bitter taste with both objective identification score (r=0.52) and intensity of the lower concentration bitter strip (r=0.51) (Supplementary Fig 3).

## Discussion

Being able to set a normal baseline of gustatory function for healthy people using a simple taste test that can be reliably self-administered at home is of immense value when looking ahead towards monitoring taste status in healthy, sick, and recovering individuals. Indeed, as the COVID-19 pandemic brought the research of smell and taste to the forefront, new quick tests of sense of smell have been introduced, for example, SCENTinel and ArOMa-T [29, 30], while quick taste testing still remains less common.

The current study focused on a taste screening test. Overall, the home and the lab groups had similar taste identification scores of the strips. High concentrations of sweet and sour obtained the highest correct identifications in both settings. Low concentrations of sour and spicy were poorly identified by the participants in this study, with even lower success for low concentrations of spicy for the home group. The present findings with a healthy population of young adults suggest that low concentrations of sour, spicy, and perhaps bitter strips are probably not suitable for the fast monitoring of gustatory function in the clinic. A possible explanation for some correlation with the bitter strips scores is the wider variation in bitterness perception, while in other taste modalities, most of the healthy participants had similar (high) scores.

Interestingly, analysis of correlations between different scores for the same individual demonstrated the highest correlation between self-reported smell and self-reported taste. This correlation is over 0.7 in the lab setting. Since in Hebrew the same word is used for “taste” and “flavor”, the confusion may be due to the dominant role olfaction plays a in the perception of flavors [31]. However, subjective report on individual taste modalities (such as bitter, sweet, salty) also had very low correlation with objective measures for that taste modality. This result can be partially due to the rather crude scales that were used here: the self-rating scales were Likert 1-7 scales and the objective identification scores were 0-2. A wider range of tastant concentrations and VAS scales may be used in future studies. The ID and mean intensity were correlated, while mean pleasantness scores had very weak negative correlations with mean intensity. The overall self-reported taste function had low or negligible correlation with the measured outcomes (identification, intensity, and pleasantness) of taste strip tasting. This further supports the notion that self-reported taste/flavor is affected by sensations other than the taste of basic taste modalities. Despite its rather small size (n=128) compared with other studies [26], the current study provides a useful initial baseline reference in a young healthy Israeli population. This can be further expanded towards standardizing a simple tool to track gustatory ability along with olfactory ability tests in a home setting [32], which is important for different impairments, for example, diabetes mellitus, head trauma, and aging [33, 34].

## Conclusion

Healthy participants reported self-assessment of taste/flavor and smell ability and then tasted paper strips impregnated with a high and low concentration of sweet, sour, salty, bitter, and of spicy (capsaicin). One group performed the test, which consisted of identification of the taste and rating of intensity and pleasantness, in the sensory lab, assisted by the researcher. The home group performed the same test in an unassisted manner, at the location and time of their choice. The results were similar in the lab and at home. The sweet and salty taste strips impregnated with 0.4 g/ml sucrose and 0.25 g/ml sodium chloride concentrations were easily identified by healthy Israeli population and provide a baseline against which taste tests can be compared in future clinical applications. Self-reported taste/flavor ability correlated with selfreported smell. Identity, intensity, or pleasantness scores from the psychophysical test did not correlate with self-reported taste, except some correlation between self-reported bitter taste and bitter taste intensity. This study illustrates the importance and practicality of simple gustatory tests in addition to self-assessment.

## Data availability

The datasets generated during and/or analyzed during the current study are available from the corresponding author on reasonable request.

## Contributions

M.Y.N, T.H., A.W.: Devising methodologies; A.W: Manufacturing strips; T.G: Participant recruitment, Examinations performed; T.G., A,O.: Analysis of data and statistics; M.Y.N, T.H, A.A., A.O, A.W, M.Y.P, T.G: Review and Editing.

