## Supplementary Figures for "Subjective assessment and taste strips testing of gustatory function, at home, and in the lab"

Supplementary material:

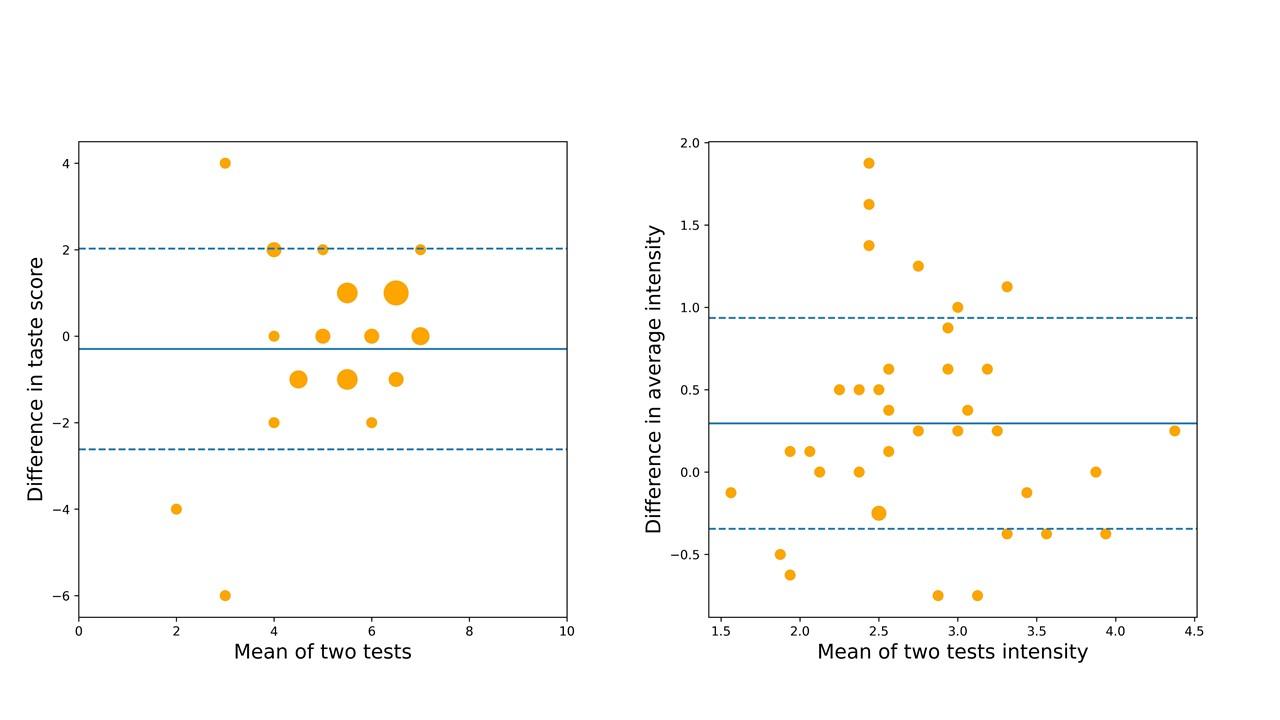

**Supplementary Fig 1.** Panel A. Bland-Altman plot. Test-retest consistency for taste identifications scores of the home group, on the Y axis is the difference between the score in the first test subtracted by the second, while the X axis indicates the mean of taste score between the two test repetitions. Mean of difference between the two tests taste score colored in blue, with one standard deviation from the mean shown in dashed line. Panel B.  indicates the consistency regarding the test-retest mean of intensity scores averaged over taste strips. Dashed line indicates one standard deviation over and under the mean of all averaged scores colored in blue.

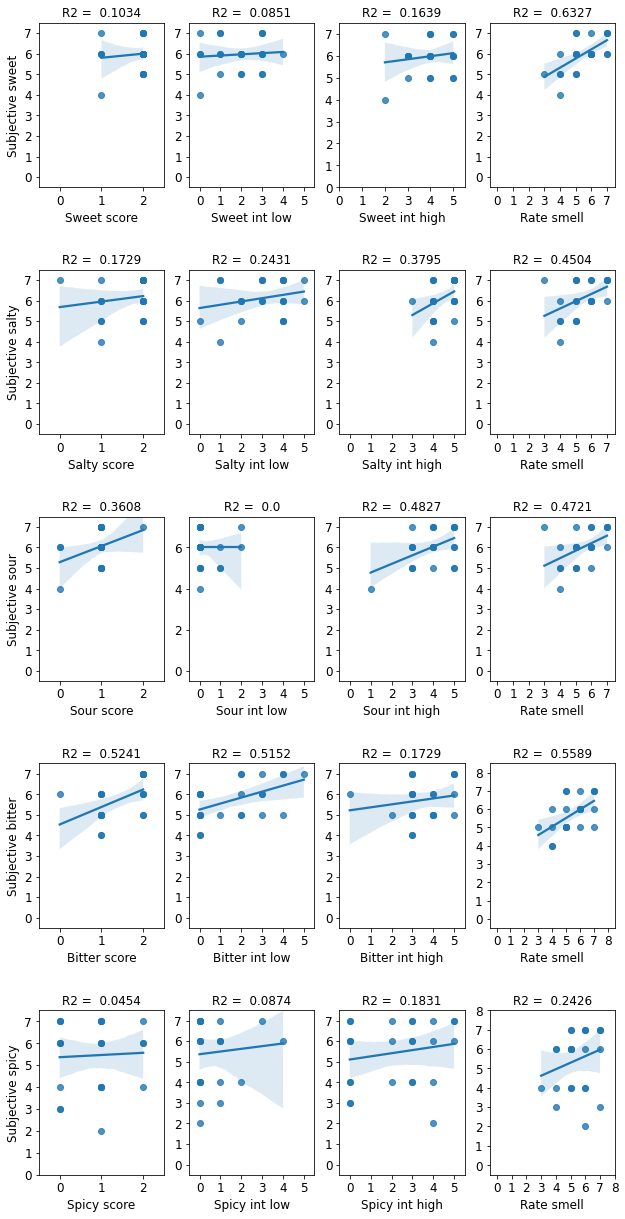

**Supplementary Fig 2.** Correlations of participants in the third group (n=25) regarding in lab scores, between each taste modality self-rating and taste strip scores, at low and high intensity.

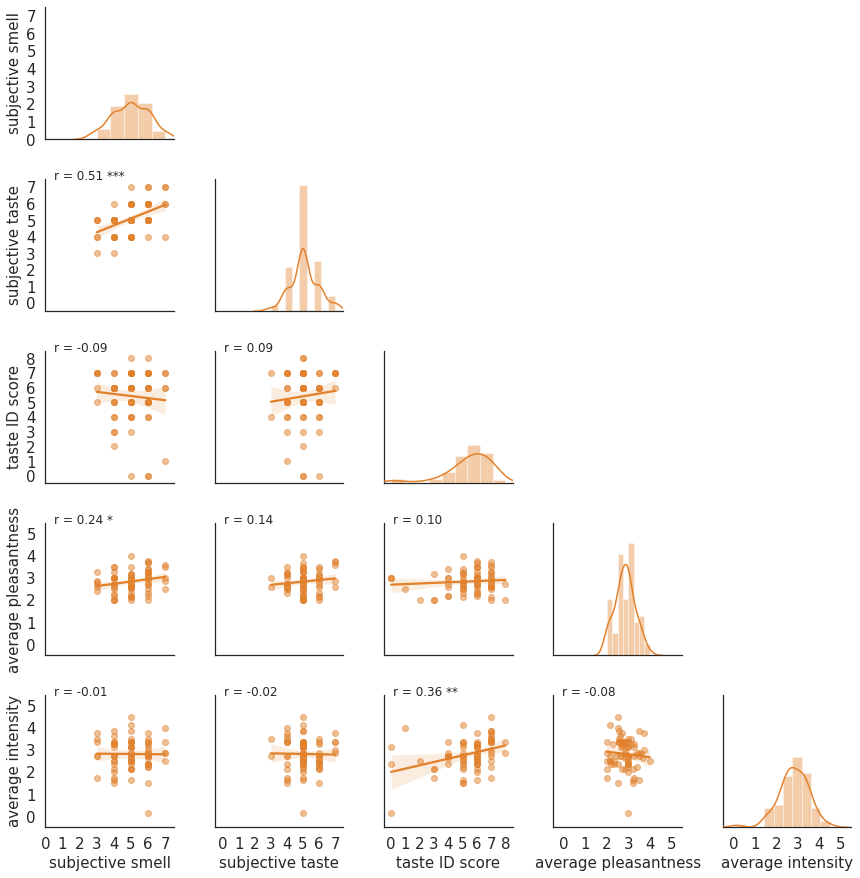

**Supplementary Fig.** 3 Correlations between the self-rated abilities to smell and taste (subjective smell and taste), taste identification score (taste ID score), and average across all of the strips regarding intensity and pleasantness ratings individually (average pleasantness and intensity), represented by scatter plots including Pearson’s R correlation coefficient at the top left of each panel, and regression line. Histograms the diagonal panels of the figure present the frequency of scores, with the x axis indicating the score and the y indicating the count of participants. Shown for home group
